# Engineered Nanotopographies Induce Transient Openings in the Nuclear Membrane

**DOI:** 10.1101/2024.08.13.605467

**Authors:** Einollah Sarikhani, Vrund Patel, Zhi Li, Dhivya Pushpa Meganathan, Keivan Rahmani, Leah Sadr, Ryan Hosseini, Diether Visda, Shivani Shukla, Hamed Naghsh-Nilchi, Adarsh Balaji, Gillian McMahon, Shaoming Chen, Johannes Schöneberg, Colleen A. McHugh, Lingyan Shi, Zeinab Jahed

## Abstract

Materials with engineered nano-scale surface topographies, such as nanopillars, nanoneedles, and nanowires, mimic natural structures like viral spike proteins, enabling them to bypass biological barriers like the plasma membrane. These properties have led to applications in nanoelectronics for intracellular sensing and drug delivery platforms, some of which are already in clinical trials. Here, we present evidence that nanotopographic materials can induce transient openings in the nuclear membranes of various cell types without penetrating the cells, breaching the nucleo-cytoplasmic barrier and allowing uncontrolled molecular exchange across the nuclear membrane. These openings, induced by nanoscale curvature, are temporary and repaired through ESCRT-mediated mechanisms. Our findings suggest a potential for nano topographic materials for direct nuclear sensing and delivery, holding promise for improving the delivery, efficiency, and safety of therapeutic agents to the nucleus.

## Introduction

Materials with engineered nano-scale surface topography are showing great promise in modulating cell-material interactions by providing unique physical cues that cells respond to. These physical cues effectively guide the interaction between materials and biological systems.^1–5^. Mimicking natural structures like viral spike proteins, nanotopographic features such as nanopillars^6^, nanoneedles^1,2^, nanotubes^7–9^ and nanowires can bypass biological barriers like the plasma membrane of cells.^10^ Due to these properties, applications of nanotopographic materials range from nanoelectronics for intracellular sensing,^4,11–14^ to both *ex vivo* and *in vivo* drug delivery platforms^1,15–18^, some of which are already in clinical trials or commercially available. Notable examples include planar patches with 3D nanostructures such as nanopillars and nanoneedles which are used for drug delivery^1,16,19^, nanopillar electrodes for neuron^14,20^ and cardiac electrophysiology^4,12^, and spiky nanoparticles that enhance cellular uptake and modulate drug release^10,21^ **(Figure 1A)**.

**Figure 1.**
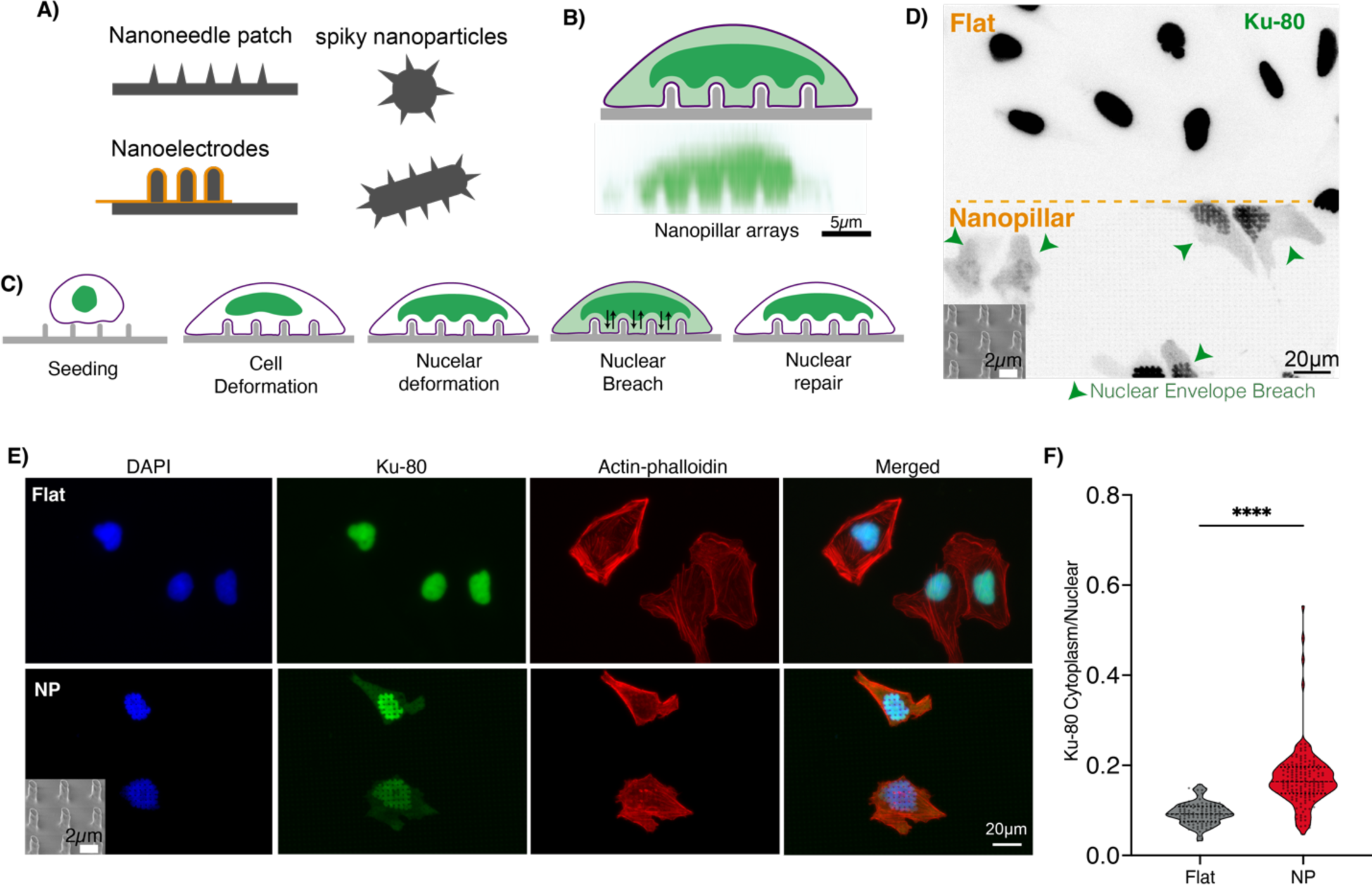
Engineered Nanotopographies for Breaching the Nuclear Membrane. A) Graphical illustration of natural mimicking nanostructures including nanoneedle patches, nanoelectrodes and spiky nanoparticles B) Schematic of nanopillars arrays inducing nuclear deformation and side-view confocal microscopy images of nanopillars with U2OS cells stained with Ku-80 antibody. The representative image of cells demonstrates the ability of nanopillars to induce nanoscale curvature on the nucleus. C) A schematic illustrating the sequence of cell-nanopillar interactions leading to cellular deformation, nuclear deformation, nuclear envelope breach, and the subsequent repair process. D) Fluorescent microscopy images of cells on flat versus nanopillars immunostained with Ku-80 antibody demonstrates the diffusion of Ku-80 to the cytoplasm on nanopillar areas due to the nuclear envelope breaching. E) Immunofluorescent images compare U2OS cells on flat and nanopillar surfaces, stained with DAPI (blue), Ku-80 (green), and Actin-phalloidin (red), demonstrating altered Ku-80 distribution on nanopillar interfaces. 30-degree tilted angle SEM images depict the distinctive nanopillar arrays (pitch=3.55± 0.06 µm, height= 3.1± 0.1µm, diameter= 0.88± 0.03 µm, Mean± S.D). F) Quantification of the cytoplasmic to nuclear Ku-80 ratio indicates higher cytoplasmic Ku-80 presence on nanopillar substrates (0.169 ±0.005, n=148) compared to flat controls (0.093±0.002, n=102)

Numerous studies have examined the interactions of nanotopographic materials with the plasma membrane, demonstrating that the geometry of these structures can induce plasma membrane remodeling, penetration, and increased endocytosis^3,22,23^. However, their interactions with other cellular organelles remain less understood. Recent studies have shown that nanotopography can deform cellular organelles such as the nucleus^6^ resulting in the remodeling of the nuclear envelope.^2,24^

In this work, we present the first evidence that nanotopographic materials can breach the nucleo-cytoplasmic barrier by inducing transient openings in nuclear membranes without penetrating cells, leading to a temporary and uncontrolled exchange of molecules across the nuclear membranes. We have found that these openings are induced by the nanoscale curvature on the nuclear membrane, which can be controlled by altering the dimensions of nanopillars and the duration of their interaction with cells. Additionally, we demonstrate that the nuclear membranes of various cell types can be breached using these materials, and that these breaches are temporary and repaired through ESCRT-mediated mechanisms.

Our discoveries pave the way for innovative methods of direct nuclear sensing and delivery using nanotopographic materials. We propose that by controlling the timing and duration of nuclear membrane openings and repairs using rationally designed nanotopographic materials, molecules can be precisely delivered directly to cell nuclei, potentially revolutionizing targeted therapeutic interventions.

## Results and Discussion

### Nanotopography-induced curvature triggers nuclear membrane breaches

The nanopillars were fabricated using a lithography process followed by both dry and wet etching. Detailed fabrication procedures are provided in our previous paper and the materials and method section.^25^ Previous research showed that nanostructures can induce nanoscale curvature in plasma and nuclear membranes.^26–28^

(**Figure S1**) exhibit both positive and negative curvature in their nuclear membranes (**Figure 1B**). Additionally, nanoscale indentation of the nuclear membrane by atomic force microscopy (AFM) has been shown to rupture cell nuclei.^29^ Based on this, we hypothesized that the curvature induced by nanotopography during cell adhesion and spreading could lead to breaching of nuclear membranes, resulting in uncontrolled exchange between the nucleus and cytoplasm, as illustrated schematically in **Figure 1C**.

To test this, we stained cells with a nuclear rupture reporter, Ku-80; a protein typically localized to the nucleus in most cell types that diffuses into the cytoplasm upon a breach in the nuclear membranes.^30–32^ We found that several cells (U2OS, HeLa, HEK-293, siLMNA iPSC-CM) on nanopillars exhibited leakage of Ku-80 into the cytoplasm, indicating nuclear membrane openings (**Figure 1D**). However, the plasma membranes of these cells remain intact, as evidenced by the absence of Ku-80 diffusion outside the cytoplasm(**Figure S2**). Notably, no such leakage was observed on the smooth regions of the same platform that lacked nanopillars (**Figure 1D**). To quantify nuclear opening events, we performed immunostaining to delineate nuclear and cytoplasmic boundaries using DAPI and Actin, respectively (**Figure 1E**), and calculated the cytoplasmic-to-nuclear ratio of Ku-80. A significant increase in this ratio was identified as a nuclear opening event, as described in the image analysis of methods section. Our analysis revealed that U2OS cells on nanopillar substrates exhibited a significant increase and a much broader range in the cytoplasmic-to-nuclear ratio of Ku-80 (0.169 ±0.005) compared to those on flat substrates (0.093±0.002), indicating a high incidence of nuclear membrane breaches on nanopillars (**Figure 1F**). Similarly, HeLa cells exhibit similar trend with cytoplasmic-to-nuclear ratio of Ku-80 on nanopillar (0.129 ±0.018) showing significantly higher value than flat substrate (0.023±0.002), indicating the breaching of nuclear envelope is not limited to U2OS cells. (**Figure S2**)

These results demonstrate that cell-nano-topography interaction can induce openings in the nuclear membrane without any other external stimulus, likely due to the nanoscale curvature induced by nanotopographies. Next, to assess whether cells with diverse structural properties would experience such nuclear membrane breaches on nanopillars, we subsequently cultured a variety of cell types on them.

### Various cell types exhibit curvature-induced nuclear membrane openings on nanotopography

To explore the versatility of nanotopography in inducing nuclear membrane openings, we investigated a range of cell types with diverse structural and morphological properties, including epithelial-like cells (U2OS, HeLa, HEK-293), stem cell-derived cardiomyocytes (siLMNA iPSC-CM), and NIH-3T3 fibroblast cells (**Figure 2A-C**). All tested cell types exhibited curvature-induced nuclear membrane openings on our nanopillar platforms. In epithelial cells and iPSC-CMs, nuclear membrane openings were evident from the mislocalization of Ku-80 from the nucleus to the cytoplasm (**Figure 2A,2B**). Conversely, in NIH-3T3 cells we found that Ku-80 was predominantly located in the cytoplasm under non-breached conditions, consistent with previous studies. Upon nuclear membrane breaching induced by nanopillars some Ku-80 localized to the nucleus.^30^ These results not only indicate the versatility of nanotopographies to induce nuclear membrane openings in various cell types, but also highlights bidirectional transfer of proteins between the nucleus and the cytoplasm during such breaches.

**Figure 2.**
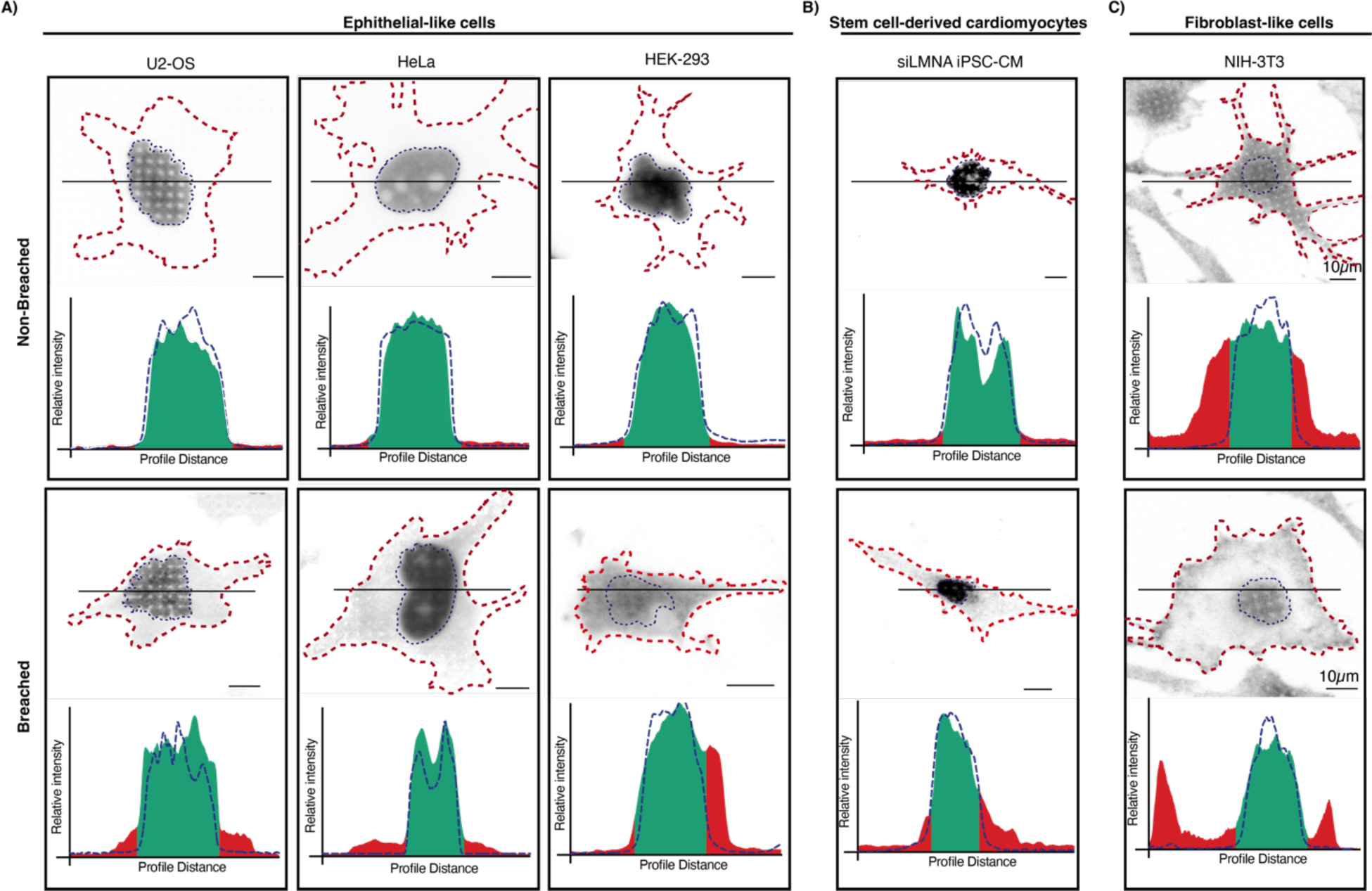
Analysis of Nanotopography-Induced Nuclear Envelope Openings Across Various Cell Types. A) Top: fluorescent image of nuclei of epithelial cells (U2OS, HeLa, HEK293) with a Ku-80 stain inside the nucleus, Bottom: cells with breached nuclear membranes as indicated by Ku-80 mislocalization to the cytoplasm. The intensity profiles of Ku-80 are shown. The red dotted line represents the cell boundary, and the blue dotted line indicates the nuclear boundary. In the intensity profiles, the red line represents the cytoplasmic region, the green line indicates the nuclear area, and the blue dotted line shows the nuclear intensity based on the DAPI channel. B) Stem cell-derived-like cells (siLMNA iPSC-CM) displaying non-breached (top) and breached (bottom) nuclear membranes on nanopillars. C) Fibroblast cells (NIH-3T3) demonstrate distinct Ku-80 translocation in the cytoplasm in cells (Top). Upon Nanopillar-induced breach of the nuclear membranes, Ku-80 translocated to the nucleus as depicted by intensity profiling (bottom).

### Nuclear membrane breaches can be controlled by nanopillar geometry and the duration of nanopillar-cell interactions

Having shown that nanotopographies can induce nuclear membrane openings in various cell types, we set out to investigate the temporal dynamics of nuclear membrane openings on nanopillars for one cell type (U2OS). In separate experiments, we quantified the number of breached nuclear membranes on nanopillars at 1, 5, and 8 hrs after seeding (**Figure S3**). Our results show that a small percentage of cells exhibit openings as early as 1-hour of cell-nanopillar interactions, with breached nuclear membranes increasing from 6.5% to 20.5% during the first 5 hours. No significant changes were observed in this percentage between 5 and 8 hours. **(Figure 3A**). To understand the mechanisms, we performed confocal microscopy to quantify the indentation depth and maximum curvature of the nuclear membrane at various durations of cell-nanopillar interactions **(Figure 3B**). We observed a significant increase in the indentation depth from 1.57± 0.19 µm at 1 hr, increasing sharply to 2.53± 0.24 µm at 5 hrs. Interestingly no significant increase was observed in the indentation depth between 5 and 8 hours, similar to non-significant breach incident difference between 5 and 8 hrs (**Figure 3B**). These results suggest that deeper indentation correlates with an increased likelihood of nuclear membrane opening (**Figure 3C**). This is further evidenced by the lack of any significant increase in the percentage of cells with nuclear membrane breaches between 5 and 8 hours, as no significant increase is observed in their indentation depths (**Figure 3C**). Additionally, the normalized maximum curvature values show a consistent increase from 0.65±0.15 µm at 1 hr to 1.59±0.41 µm and 2.34±0.84 µm at 8 hrs suggesting that prolonged exposure to nanopillars increases the curvature and deformation of nuclear membrane on nanopillars which leads to more frequent nuclear membrane breaches. (**Figure 3D**)

**Figure 3.**
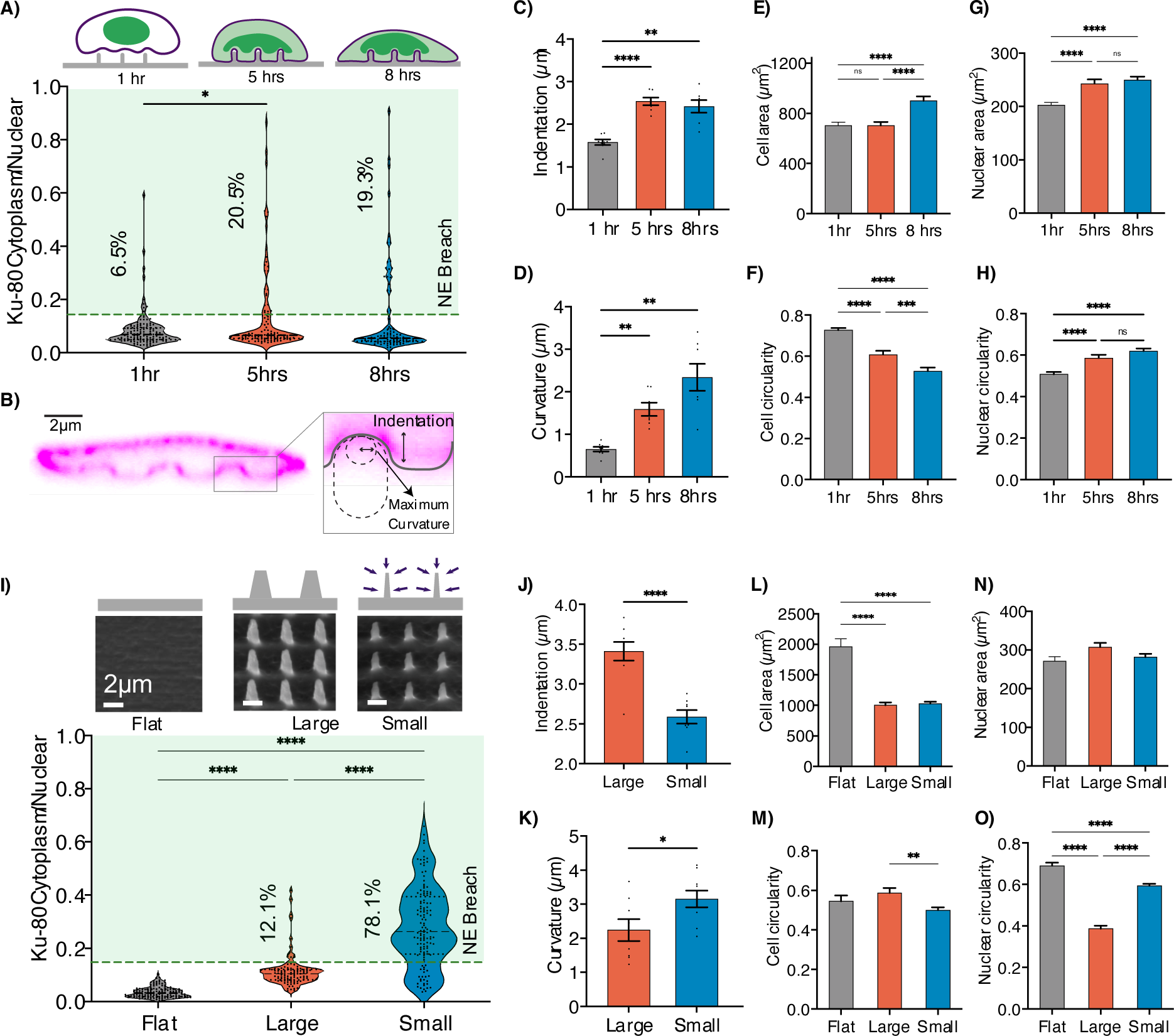
Temporal and Size-Dependent Effects of Nanotopography-Induced Nuclear Membrane Openings. **A**) The schematic and quantitative analysis highlights a significant increase in the cytoplasmic to nuclear Ku-80 ratio from 1 to 5 hours. A cytoplasmic/ nuclear Ku-80 ratio threshold of 0.16% is established to identify nuclear membrane openings. Data indicate that 6.5% of cells exhibit nuclear membrane breaches at the 1-hr point, with the proportion of breached cells increasing to 20.5% and 19.3% at 5 and 8 hours, respectively. (n>92). **B)** The side-view immunofluorescent image of Lamin A/C on nanopillars showing the curvature and indentation depth of the nucleus due to the nanopillars. **C)** The indentation depth of the nuclear envelope at 1, 5, and 8 hrs post seeding illustrates a significant increase from 1.57± 0.19 µm to 2.53± 0.24 µm and 2.41±0.39 µm (n>8 cells per condition. Each value represents the mean values of indentation induced by each nanopillar). **D)** the normalized maximum curvature of the nuclear envelope, showing an upward trend from 1 hour to 8 hrs. (n>8 cells per condition. Each value represents the mean values of curvature induced by each nanopillar). **E)** Cells have higher area from 5 to 8 hrs indicating the cellular spreading while the breach incidence remains unchanged between 5 and 8 hrs indicting the breaching likely is not dependent on cellular area. **F)** Cell circularity measurements suggest an initial round morphology at 1 hr, which decreases over time, exhibiting higher cell deformation. **G)** The nucleus area parameter shows a full spread of the nucleus by the 5-hr time point with no further changes through to the 8-hr time point. **H)** The nuclear circularity increases by 5 hrs and then stabilizes up to the 8-hr time point. The nuclear circularity is higher after cells spread suggesting nuclear deformation while cells are in early adhesion and spreading phase. **I)** The tilted 30-degree SEM images of flat, large (height= 4.70±0.02µm, diameter= 1.30±0.01µm) and small nanopillars (height= 3.18±0.03µm, diameter= 0.51±0.03µm), schematic of isotropic etching of nanopillars and cytoplasmic/nuclear Ku-80 ratio, showing notably higher presence of Ku-80 in the cytoplasm on small nanopillars compared to large nanopillars and flat substrate. Small nanopillars exhibit the highest incidence of nuclear envelope breach at 78.1%, while large nanopillars demonstrate a significantly lower breach rate of 12.1%. (n≥98). **J)** Indentation depth of nuclear envelope associated with large and small pillars, deeper indentation for larger pillars 3.41± 0.34 µm compared to small pillars 2.58± 0.24 µm (n>8) **K)** The normalized maximum curvature of nuclear envelope where the small nanopillars induce higher curvature 3.15± 0.73 µm compared to large pillars 2.24± 0.90µm. (n>8). **L)** Analysis of cell area suggests that cells on both large and small nanopillars exhibit a reduced spreading compared to flat surfaces. **M)** Cell circularity measurements indicate greater cell deformation on large nanopillars when compared to flat and small nanopillar substrates. **N)** Nuclear area analysis detects no significant difference in nuclear spreading across different substrate topographies **O)** The nucleus circularity between small, large and flat indicating the highest nuclear deformations on small nanopillars.

Furthermore, we hypothesized that the spreading of cells and nuclei on nanopillars leads to increased nuclear membrane breaches. To explore this, we quantified cell and nuclear morphologies, specifically spreading (area) and shape (circularity), at 1, 5, and 8 hrs (**Figure 3E-H**. Although cell area remained largely unchanged from 1 to 5 hours (**Figure 3E**), cell shape changed significantly (**Figure 3F**), indicating cell deformation around the nanopillars during this period. More importantly, significant changes in nuclear area (**Figure 3G**) and shape (**Figure 3H**) were observed between 1 and 5 hrs, aligning with the rise in nuclear membrane breaches. These metrics remained unchanged from 5 to 8 hrs (**Figure 3G,3H**), underscoring that nuclear spreading and deformation is key to nuclear membrane breach on nanopillars. Interestingly, although cell spreading area and shape continued to change significantly from 5 to 8 hrs (**Figure 3E, 3F**), these alterations did not correlate with any further increase in nuclear membrane breaching. This finding indicates that changes in nuclear dimensions and shape, rather than those of the cell, are the primary drivers behind the nuclear membrane breaching on nanopillars.

Together these results show that the incidence of nuclear membrane breaching can be controlled by adjusting the duration of interactions between the nanopillars and cells.

To investigate whether the size of nanopillars could influence the frequency of nuclear membrane breaches, we employed isotropic wet etching to reduce nanopillar dimensions, keeping other factors constant. (**Figure 3I)** We compared the percentage of cells with nuclear membrane openings on large (height= 4.7±0.02 µm, diameter= 1.30±0.01µm) and small pillars (height= 3.18±0.033 µm, diameter= 0.515±0.03 µm). (**Figure S4**) Interestingly we found an increase in the percentage of cells with breached nuclear membranes from 12% to 78% with nanopillar size reduction, while smooth surfaces showed none (**Figure 3I**). We found that although smaller nanopillars induce a lower indentation depth (2.58± 0.24 µm) compared to larger nanopillars (3.41± 0.34 µm) the normalized maximum curvature of the nuclear membrane is much higher on smaller nanopillars (3.15± 0.73µm) compared with larger ones (2.243± 0.90µm) (**Figures 3J, 3K**). The higher curvature associated with smaller nanopillars appears to be the critical factor in the increased frequency of nuclear envelope breaches. Despite the larger pillars causing deeper indentation, their relatively lower curvature contributes to a reduced breaching of the nuclear membrane suggesting that while indentation depth contributes to the mechanical disruption of the nuclear envelope, it is the degree of curvature that more significantly dictates the likelihood of formation of membrane openings.

To assess whether changes in cell and nuclear morphology, specifically in area and circularity, contribute to the higher incidence of nuclear membrane breaches on smaller nanopillars, we compared these aspects of cell and nuclear structure. Our analysis showed that cells on nanopillars had a smaller spreading area than those on flat surfaces, regardless of the nanopillar size, with no notable difference in cell area between small and large nanopillars. (**Figure 3L**). Additionally, we found that cell and nuclear circularity, rather than spreading area, varies between small and large nanopillars (**Figure 3L-3O**). This suggests that the distinct cell and nuclear circularity imposed by the size of the nanopillars may play a role in nuclear membrane breaching (**Figure S5**).

### The nuclear membrane is rapidly repaired after a nanotopography induced breach

We next investigated whether nuclear membrane breaches induced by nanopillars are transient.^33^ By conducting live-cell imaging of cells expressing NLS-GFP as a nuclear rupture reporter, we observed that the nuclear membrane repairs itself within approximately 1.5 hrs after the breach, as shown in **Figures 4A and 4B**. This repair process is evident from the gradual decrease in the cytoplasmic to nuclear ratio of NLS-GFP following breach, followed by its complete relocalization to the nucleus after 1.5 hrs, indicating a successful repair mechanism. Additionally, the initiation of a rupture event coincided with the localized expression of a cytoplasmic DNA-binding protein fused to mCherry, Cyclic GMP-AMP synthase (cGAS), which marks the opening site and indicates that some DNA is exposed to the cytosol at these locations^34,35^ (**Figure 4C,D**). Furthermore, cGAS remained at the breach site even after the nuclear membrane was fully repaired and NLS-GFP had relocalized to the nucleus, corroborating previous studies that have demonstrated the persistent presence of cGAS at nuclear membrane rupture sites during cancer cell migration.^36,37^ Finally, through two-photon microscopy, we show that ESCRT-III repair machinery, specifically CHMP4B, are recruited to nuclear membrane openings and aid in repairing the nuclear membrane (**Figure 4E**).^38,39^ Altogether, these results suggest that nanotopography-induced nuclear membrane openings are repairable and transient.^40^

**Figure 4.**
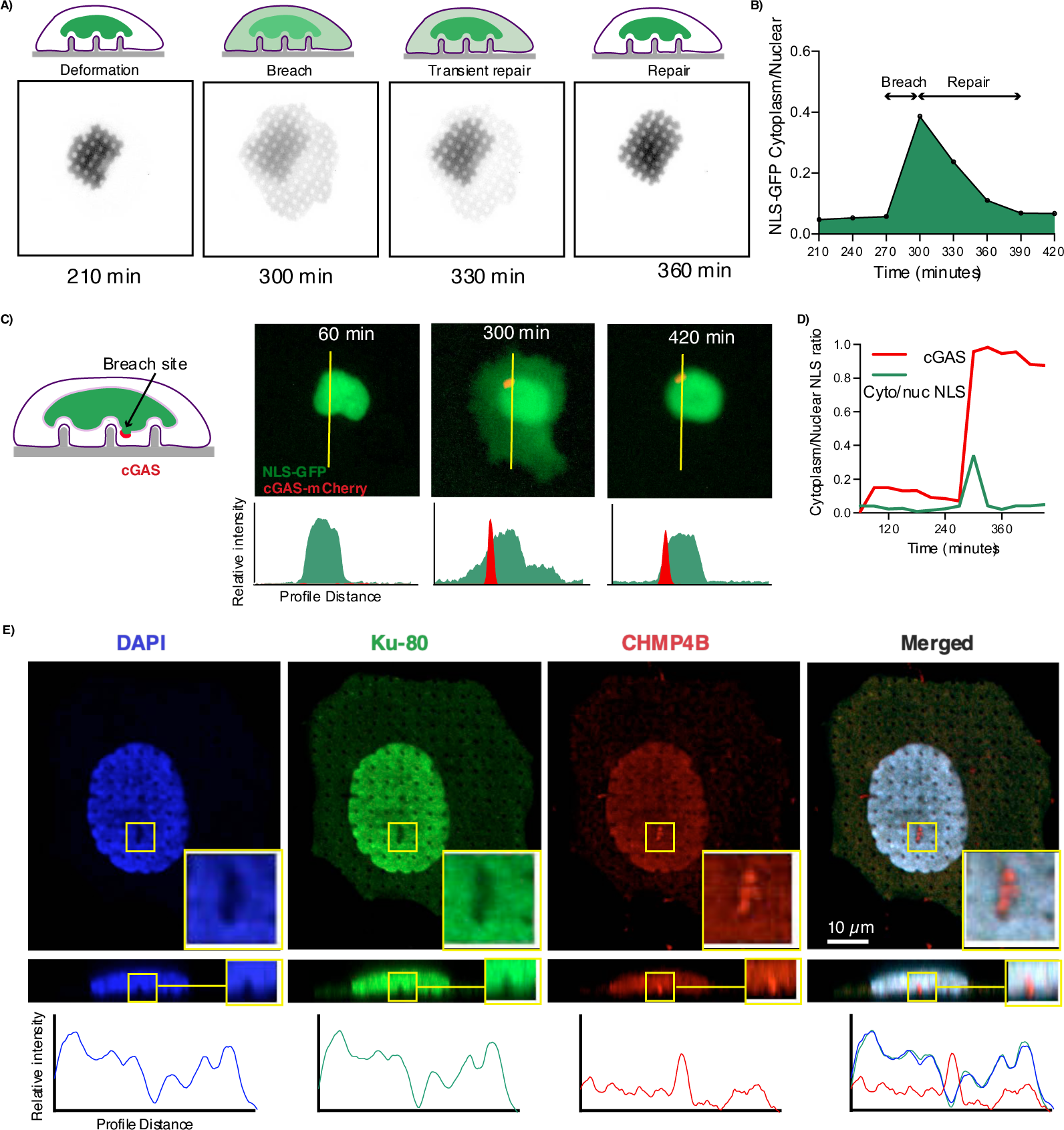
Repair of nanotopography-induced nuclear membrane breaches through ESCRT mediated mechanisms. A) Time-lapse imaging shows U2OS cells expressing NLS-GFP undergoing nuclear membrane breach and repair. Notably, after 90 minutes, cells display signs of breaching with subsequent repair mechanisms relocalizing NLS-GFP to the nucleus after 2.5 hrs. U2OS cells. B) The cytoplasmic and nuclear NLS-GFP signal ratio illustrates the decreasing ratio after the rupture incident suggesting the repair mechanism. C) Overall schematic and time lapse of images of U2OS cells expressing NLS-GFP and cGAS-mCherry showing membrane breach and repair. The cGAS-mCherry binds to the DNA exposed to the cytoplasm upon breaching. Cells showing breaching after 5 hrs by mislocalization of NLS to the cytoplasm and showing the cGAS signals. Additionally, cells repair after 7 hrs while the cGAS remains in the breach sites which is indicated by the profile analysis of cGAS and NLS over the yellow line. Representative images of U2OS cells transfected with NLS-GP and cGAS-mChery showing the site of the membrane breach on nanopillars induced by the curvature. D) Profile intensity of the NLS and cGAS shows increased cytoplasmic to nuclear NLS ratio follows by sudden increase in the cGAS signals which confirms the presence of cGAS and NLS upon membrane opening. Additionally, after 7 hrs Ku-80 relocalized to the nucleus while cGAS signal is still present. E) Immunofluorescence microscopy of U2OS cells, viewed both from the top and the side. The nuclei are stained with blue DAPI stain; Ku-80, which signals nuclear envelope breach in green; and CHMP4B ESCRT-III protein which is implicated in nuclear membrane repair in red. The highlighted yellow region within the images shows the area of nuclear envelope disruption, characterized by gaps in the DAPI and Ku-80 where CHMP4B is localized, suggesting active repair processes. Side-view images further emphasize CHMP4B’s presence at the sites of nuclear breach and repair on the nanopillars.

## Conclusions and future directions

The nuclear membrane serves as the main barrier regulating selective molecular exchange between the cytoplasm and nucleoplasm. Overcoming this barrier is crucial for delivery of cargo into the nucleus of cells and for applications in intra-nuclear sensing. Here we show that nanotopography can temporarily breach the nuclear membrane of many cell types. We further showed that by tuning their geometries and duration of interaction with cells, the nanotopographic features can control the incidence of nuclear membrane breaches.

Our future work will concentrate on the direct delivery of various biomolecules into the nucleus of cells employing nanotopography. With capabilities to directly observe nuclear opening and repair events, we can link these occurrences to nuclear delivery events. This will enable us to intelligently design our nanotopographic features to achieve nuclear delivery in various cell types. The results presented in this paper increase our confidence that we can apply this method in diverse cell types for various therapeutic applications.

## Materials and Methods

### Nanopillar Fabrication

Nanopillar chips were fabricated on a 4-inch fused quartz wafer (Wafer Pro) and underwent RCA cleaning using both SC1 and SC3 solutions, followed by a spin rinse and drying using a Spin Rinse Dryer (MEI). AZ 1512 photoresist (EMD Performance Materials) was spin coated at 4000 rpm for 45 seconds, accelerating at 12000 rpm during a three-step procedure, achieving a photoresist thickness of ∼ 1.2 µm. Our custom design file (.gds) was used to pattern the wafers, which were then exposed using the Heidelberg MLA system at a wavelength of 375 nm and a dosage of 300. The substrate developed using AZ400 developer (AZ Electronic Materials) for 30 seconds. Chromium (Cr) with a purity of 99.998% was deposited onto the patterned substrates via a Temescal Ebeam evaporator. A lift-off process was carried out using RR41, acetone, and isopropyl alcohol (IPA) in sequence. The substrates dry etched for 50 minutes in an Oxford Plasmalab 80 using Reactive Ion Etching (RIE) system using argon (Ar) at 35 sccm and chlorotrifluoromethane (ChF3) at 25 sccm, under conditions of 50 millitorr and 200 watts.

Wafer wet etched for 10 minutes using a chromium etchant (Transene Company Inc.) followed by wet etching in buffered oxide etch (BOE) 20:1 to remove the exposed quartz. The final step involved dicing the wafers into chips measuring 1 cm x 1 cm for further functionalization.

### Nanopillar Preparation

For Gelatin NP chips were sterilized by rinsing with 70% ethanol, followed by deionized water, and air-dried. The chips were UVO treated for 10 minutes and incubated with 100 µg/mL poly-L-lysine (PLL) solution. PLL was applied by inverting the chip onto a 50 µL droplet on parafilm, with the nanopillar side contacting the droplet, and incubated at room temperature for 30 minutes. The chip was then washed 3 times with PBS. 0.5% glutaraldehyde solution added for 30 minutes to crosslink the proteins and washed with PBS. Finally, a pre-warmed 0.1% gelatin solution was applied, and the chip was incubated for another 30 minutes at room temperature.

To coat the nanopillar chips with FN for iPSC-CM and NIH-3T3 cells, we sterilized the chips and rinsed with 70% ethanol followed by deionized water, allowing them to air dry after each rinse. We have applied 50 µL of 50 µg/mL FN and incubated the chips in incubator at 37°C for at least 2 hours or alternatively overnight at 4°C.

### Cell Culture and seeding

U2OS and NIH-3T3 cells were obtained from ATCC; HEK-293 cells were a gift from Dr. Ester Kwon, and HeLa cells by Dr. Lingyan Shi. U2OS cells were cultured in McCoy’s 5A medium (ACC) supplemented with 10% fetal bovine serum (FBS, Invitrogen) and 1% penicillin-streptomycin (PenStrep, ThermoFisher Scientific). HeLa cells were maintained in Eagle’s Minimum Essential Medium (EMEM, ATCC) with the same supplements. HEK-293 cells were grown in Dulbecco’s Modified Eagle Medium (DMEM, ThermoFisher Scientific) with 1% PenStrep, and NIH-3T3 cells were cultured in DMEM supplemented with 10% FBS and 1% PenStrep. All cell lines were cultured under humidified conditions at 37°C and 5% CO2.

To seed cells onto NP chips, first we have detached cells using TrypLE ™ Express Enzyme (Gibco) and prepare a 100 µL suspension containing 50,000 cells. The NP chips placed in a 24-well plate and rinse it three times with PBS. We have added the droplet of cells and mix thoroughly to ensure uniform distribution of cells. After 10 minutes of initial adhesion, we add 1 mL of growth media and incubate cells at 37°C for further analysis.

### Plasmid transduction

Plasmid was expanded according to the Addgene agar stab protocol. Briefly, a bacterial colony was collected from the stab and streaked onto an LB agar plate (Sigma Aldrich).

All plasmids were streaked from provided bacterial stabs onto agarose plates containing the respective antibiotic and incubated overnight at 37°C. Individual colonies for each plasmid were selected and grown overnight in 10mL of LB Broth incubated with the appropriate antibiotic, at 37°C with shaking. After overnight growth, plasmid DNA was extracted by miniprep with the QIAprep Spin Miniprep Kit (ID 27104) from Qiagen. Miniprep procedure was followed as recommended by the manufacturer. Purified DNA plasmids were eluted in ultrapure H 2 O and DNA concentration and purity was validated through A260 ratios on a NanoDrop Lite Spectrophotometer from ThermoFisher.

### Cell lines and transfections

U2OS cells were transfected to express nuclear rupture indicator NLS-GFP (pCMV-PV-NLS-GFP) and cGAS-mCherry (pCDH-cGAS-E225A D227A-mCherry2-EF1-Puro for live imaging. The cells were starved in serum-free minimal medium (OPTI-MEM) for 30 minutes. A complex of 1 μg of plasmid was mixed with 3 μl of Viafect (Promega) in OPTI-MEM and mixed and incubated at room temperature for 20 minutes. The media exchanged with serum-free McCoy 5a, and complex added to the cells. After 1 day transfection medium was replaced with growth media and cells recovered before live imaging.

### Immunostaining

Cells fixed using 4% paraformaldehyde (Electron Microscopy Sciences, USA), for a duration of 10 minutes at room temperature and washed with PBS to remove excess fixative. Cells were permeabilized with a 1% solution of Triton™ X-100 (Sigma-Aldrich, USA) for 10 minutes and blocked using a 2% w/v solution of Bovine Serum Albumin (BSA) (Thermo Scientific™, USA) for one hour.

Antibodies used for immunofluorescence were Rabbit anti-Ku-80 (1:200, Cell Signaling), Mouse anti-Lamin A/C (1:400, Biolegend), and Rabbit anti-CHMP4B (1:100, Novus Biologicals). Secondary antibodies conjugated to Alexa Fluor 488, 594, or 647 (Thermo Fisher Scientific) were used at 1:1,000. Samples were washed in PBS and stained with 4’,6-diamidino-2-phenylindole (DAPI) (Thermo Scientific™, USA) for 5 minutes and Alexa 594-phalloidin (Invitrogen™, USA) for 20 minutes in the dark to stain the nucleus and F-actin.

### Time lapse imaging

For live imaging, nanopillar chip was placed on a glass bottom well plate. We used Echo Revolution System with 20× Olympus objective with stage top incubator (Live Cell Instrument) for the duration of the experiment. Cells were maintained at 37°C and 5% CO2 throughout the experiment.

### SEM imaging

Scanning Electron Microscopy (SEM) was performed using a FEI Apreo FESEM system. Imaging was conducted under low vacuum conditions and operated at 5 kV voltage. To enhance the visibility of the nanopillars’ height, images were captured at a 30-45-degree tilt.

Cell samples were fixed and dehydrated for SEM imaging. Briefly, 2.5% glutaraldehyde added and incubate overnight at 4 degree and rinsed with PBS at least 3 times. Samples dehydrated through a series of ethanol washes, increasing the concentration from 30% to 100%, with each step lasting 10 minutes. Finally, sample covered with Hexamethyldisilazane (HMDS) for 15 minutes and HDMS removed and air dried overnight before SEM imaging.

### Fluorescence and two photon fluorescence microscopy

Images were collected with Echo Revolution microscope with following objectives. 20x PLAN Fluorite LWD CC Phase Ph1 NA 0.45, 40x PLAN Fluorite LWD CC Phase Ph2 NA 0.60, 60x PLAN Fluorite Water Dipping NA 1.00.

An upright laser-scanning microscope (DIY multiphoton, Olympus) with a 25x water objective (XLPLN, WMP2, 1.05 NA, Olympus) was applied for near-IR throughput. The setup included a synchronization of pulsed pump and stokes beams through a picoEmerald system (Applied Physics & Electronics), with the pump beam tunable between 780 and 990 nm, a pulse width of 5-6 picoseconds, and a repetition rate of 80 MHz, while the Stokes beam was fixed at 1031 nm with identical pulse width and repetition rate. Both beams were directed into the microscope and the transmission was collected via a high numerical aperture (NA) oil condenser (1.4NA). For the two-photon fluorescence (TPF) imaging, 781 nm and 810 nm pump beams were employed for capturing DAPI and FITC stained images, respectively. Each pixel was exposed for 12.5 microseconds, and images were averaged over three frames. The backscattered light was separated using a dual filter cube with passbands at 460 nm and 515 nm. All images were captured at a resolution of 512 × 512 pixels, and a z-step size of 1µm.

### Image analysis

ImageJ 1.53 (NIH, US) used for image analysis to quantify nuclear envelope rupture via the cytoplasmic to nuclear ratio of Ku-80 protein intensity. Images were processed to select the nuclear and cytoplasmic area using immunofluorescence staining; Actin staining provided the cellular outline, while DAPI staining defined nuclear regions. To calculate the cytoplasmic/nuclear ratio of Ku-80, intensity measurements were performed separately on cytoplasmic and nucleic regions. Initially, the DAPI channel was used to threshold and define nuclear areas, followed by measurement of Ku-80 intensity within these boundaries. Subsequently, the Actin channel outlined the entire cell for assessing total cellular Ku-80 intensity. The area and circularity of cells and nucleus calculated using analyze particle command as follow:

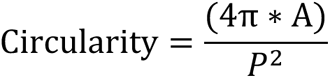

Where A shows the area and P is the indicator of perimeter.

To quantify the curvature characteristics of an ellipse relative to a circular geometry with an equivalent area, we have resliced the images from confocal imaging and reconstructed the side view of the images for fitting the ellipse. We calculated the normalized maximum curvature (κ_norm_) using imageJ for calculating the major axis (a) and minor axis (b) of ellipse.

The maximum curvature of the ellipse is defined as below, where we have assumed that *a* > *b* > 0:

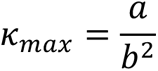

To normalize the ellipse’s curvature, we used the curvature of a circle whose area is equal to that of the ellipse. The curvature of a circle with the same area as the ellipse defined by:

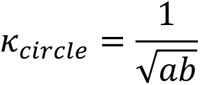

The normalized maximum curvature is then calculated by dividing the maximum curvature of the ellipse by the curvature of the equivalent circle:

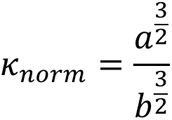

To differentiate between ruptured and non-ruptured cells based on Ku-80 intensity ratios, we utilized Gaussian Mixture Models (GMM). This statistical method models data as a combination of Gaussian distributions, each representing a different cluster within the dataset. In our analysis, we applied GMM to identify two clusters: one for ruptured cells and another for non-ruptured cells. The intersection of these distributions provided a data-driven threshold, optimizing the distinction between the two states. The GMM determined a threshold value of 0.16, which effectively separated cells with ruptured nuclei from those intact, highlighting its utility in refining the analysis without relying on arbitrary preset values.

### Statistical analysis

Unless otherwise noted, all experimental results were taken from at least three independent experiments. For comparisons, we used student’s t-tests (comparing two groups) or one-way ANOVA (for experiments with more than two groups) with post-hoc tests. All tests were performed using GraphPad Prism., * denotes p ≤0.05, ** denotes p ≤0.01, and *** denotes p ≤0.001. Unless otherwise indicated, error bars represent the standard error of the mean (SEM)

## Acknowledgements

This work was performed in part at the San Diego Nanotechnology Infrastructure (SDNI) of UCSD, a member of the National Nanotechnology Coordinated Infrastructure, which is supported by the National Science Foundation (Grant ECCS-2025752). We’d like to thank Dr. Peng Guo and the Nikon Imaging Center at UCSD for the support on microscopy experiments. This work was in part supported by Air Force Office of Scientific Research YIP award (Award Number: 311616-00001) and Cancer research coordinating committee faculty seed grant to Z.J. This work was supported by NIH R00 GM12049403 awarded to C.A.M. S.S is supported through NSF GRFP Fellowship. NIH NIBIB Award to G.M. under Award number T32EB009380. We acknowledge the support from UCSD Startup funds, NIH R01GM149976, and NIH R21NS125395 to LS.

